# Transcriptional Regulation of the Alternative *Sex Hormone-Binding Globulin* Promoter by KLF4

**DOI:** 10.1101/2023.10.13.562308

**Authors:** Warren M. Meyers

## Abstract

In most mammals the major site of sex hormone-binding globulin (SHBG) synthesis is the liver wherefrom it is secreted into the bloodstream and is the primary determinant of sex steroid access to target tissues. The minor site of SHBG synthesis is the testis and in lower mammals testicular SHBG has long been known to be synthesized and secreted by Sertoli cells. However, human testicular *SHBG* is expressed in developing germ cells from an upstream alternative promoter (altP-*SHBG*). Transcripts arising from this region comprise an alternative first exon (1A) with the resultant protein confined to the acrosomal compartment of the mature spermatozoa. I have dissected the regulatory components of the alternative *SHBG* promoter and identified motifs that are required for optimal transcriptional activity from this region. Transcriptional activity is driven by two CACCC elements that appear to be functionally redundant. The transcription factor KLF4 interacts with promoter the region spanning these elements *in vivo*. Knockdown of *Klf4* results in decreased altP-*SHBG* activity, while *Klf4* overexpression relieves the effects of knockdown. Based on their shared patterns of expression *in vivo*, I conclude that KLF4 is a transcriptional regulator of *SHBG* in male germ cells.

## 1. Introduction

The major transcription unit of the human *sex hormone-binding globulin* (*SHBG*) gene encodes the SHBG precursor that is processed in the endoplasmic reticulum of hepatocytes and secreted into the bloodstream (1,2). It comprises an ∼800 bp proximal promoter sequence that immediately flanks exon 1 (2–4). Analyses of numerous other *SHBG* transcripts have revealed that many contain alternative exon 1 sequences located from between 1.9 kb and 17 kb upstream from the major *SHBG* transcription unit expressed in the liver (Figure 1A). At least six different transcriptional start sites have been identified along the *SHBG* gene (5–8). Importantly, the only alternative *SHBG* transcript that results in a truncated SHBG isoform is the one expressed in testicular germ cells containing exon 1A, (9–11). Transcripts originating from exon 1A are spliced directly to exon 2 (5,6), bypassing exon 1 that encodes the secretion polypeptide, and resulting in a protein that accumulates within the acrosomal compartment (11). Translation of this sperm SHBG isoform is hypothesized to initiate from methionine 30 along exon 2 (11). The resultant protein has a truncated LG4 domain that lacks the N-terminal O-glycosylation site present on secreted SHBG, though possesses a steroid binding ability that is indistinguishable to SHBG in serum (9).This feature of the human *SHBG* gene is recapitulated in the testes of the humanized transgenic mice that carry an 11 kb region of the human *SHBG* locus (11K-SHBG) (9,12).

**Figure 1.**
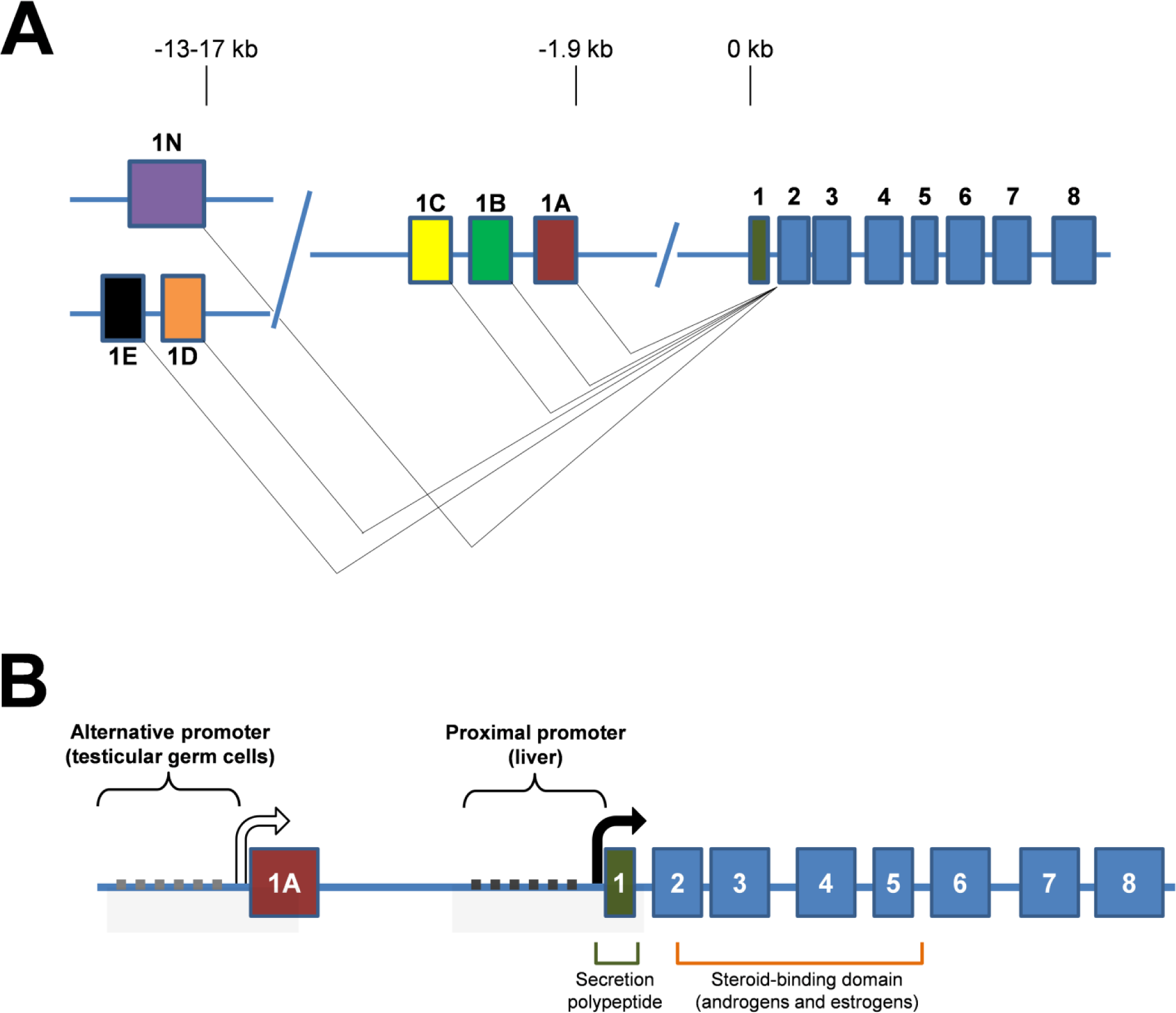
Alternative *SHBG* transcription units. A, There are at least six known alternative transcription units on the human *SHBG* gene used in different tissues. The major transcription unit used by the liver initiates from exon 1 and results in a mature protein secreted into the bloodstream (1,2). Transcription from exon 1A occurs in testicular germ cells of humans and humanized 11K-SHBG transgenic mice (3–8). The remaining transcriptional start sites are used in either the human prostate or human cancer cell lines (9,10) where their significance is unknown. B, The proximal and major alternative *SHBG* transcription units are each controlled by their own distinct promoter regions (2). Transcripts arising from exon 1A in testicular germ cells splice directly to exon 2, bypassing the secretion polypeptide yet retain the steroid-binding domain and results in a truncated intracellular isoform of SHBG. Relative positions of exons in A and B are not to scale.

Transcription of these alternative *SHBG* transcripts appear to be under the control of the promoter sequence that flanks alternative exon 1A (Figure 1B). Compared to the proximal promoter that controls *SHBG* expression in the liver, this alternative promoter produces very high transcriptional activity in the immortalized mouse spermatocyte cell line (GC-2) within the context of a luciferase reporter assay (11), mimicking what is observed *in vivo*. I first interrogated the alternative promoter sequence that drives *SHBG* expression in the testis for known regulatory elements or motifs. Presumed elements of importance were tested by mutagenesis to assess their impact in a luciferase reporter gene assay, as described previously (11). I then considered factors that interact with these elements and if they shared patterns of expression with *SHBG* during spermatogenesis *in vivo*.

## 2. Results

### 2.1 High transcriptional activity from the alternative *SHBG* promoter in GC-2 cells is dependent on intact CACCC elements

The -336/+28 bp alternative *SHBG* promoter (altP-*SHBG*) was cloned into the pGL3 luciferase reporter plasmid. When transfected into mouse GC-2 cell line the reporter activity of altP-*SHBG* was over 150-fold above background (*P*<0.0001) while the construct containing the proximal *SHBG* promoter used in hepatocytes failed to produce significant activity above background (Figure 2). I then asked what features inherent within the alternative *SHBG* promoter contributed to its basal activity in GC-2 cells. Sequence analysis of this region revealed several motifs and features of interest (Figure 3A). The -336/+28 region has 65.5% guanine-cytosine (GC) content and is rich in 5’-cytosine-phosphate-guanine-3’ (CpG) sites with 23 CpG sites on the sense strand and 22 on the antisense. This yielded a CpG frequency of ∼14.6% on the sense strand and therefore falls within the accepted definitions of a CpG island (15,16). CpG Islands locator tool confirmed this finding by scoring a CpG island from -336 to -55 along this region with an observed/expected ratio of CpG sites at >0.6. Additionally, three candidate regulatory elements were identified. Two identical CACCC (100% consensus) elements located at -198/- 189 and -111/-102 and one partial (50% consensus) cAMP response element (CRE) located at - 180/-173 (17). Sequence analyses also identified partial SPZ1 binding sites along this region as has been reported earlier (11). However, since the SPZ1 motif hits from different analysis programs did not all map to the same location, they were not pursued further.

**Figure 2.**
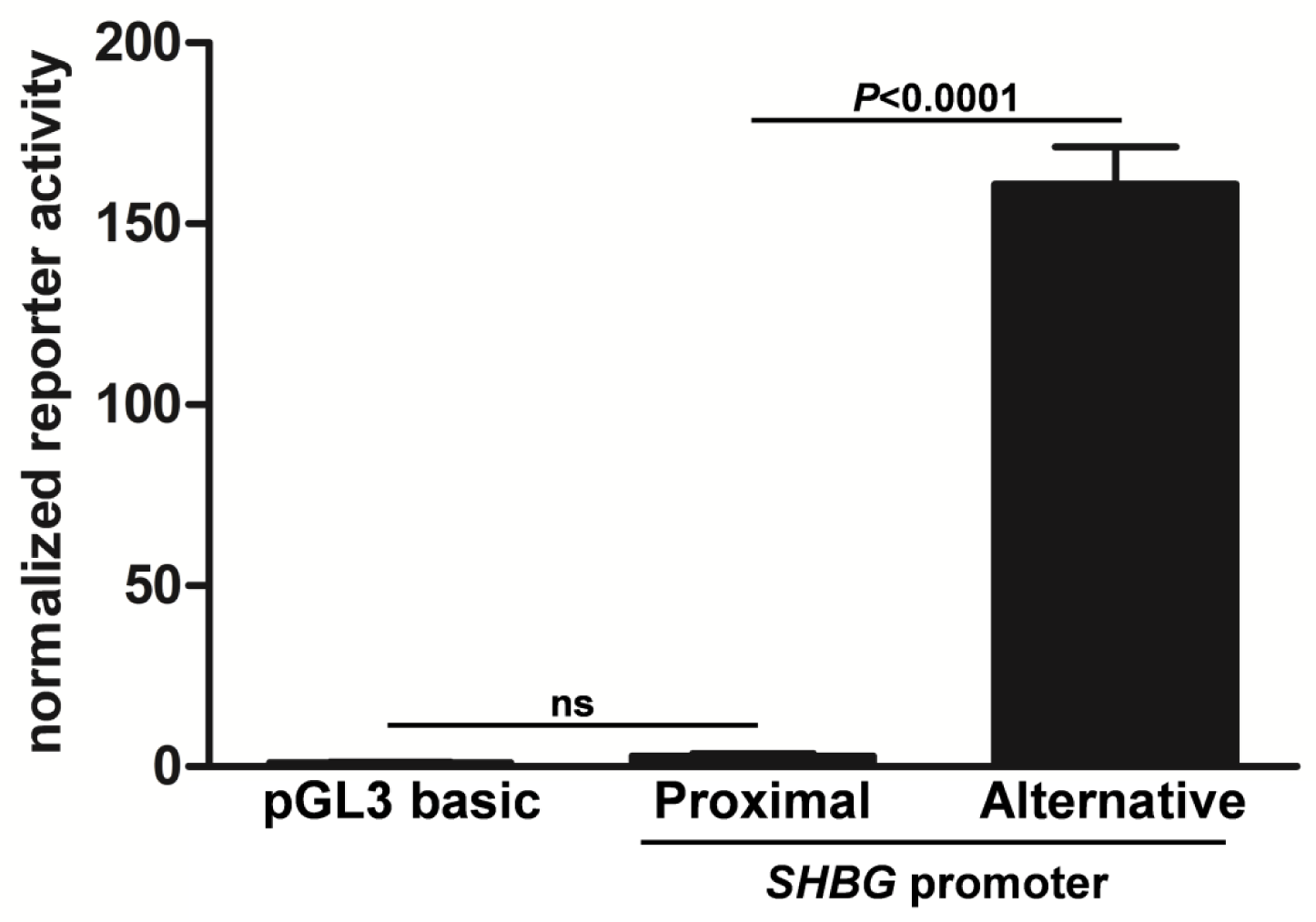
High transcriptional activity of the human alternative SHBG promoter in GC-2 cells. The basal luciferase reporter activities of the proximal (-266/+366) and alternative (-336/+28) *SHBG* promoters were assessed in GC-2 cells. Relative luminescence counts are normalized to a promoterless pGL3 basic reporter vector that was assayed in parallel. Data points are means ± SD of triplicate measurements. *P*-values and statistical significance were determined by one-way ANOVA.

**Figure 3.**
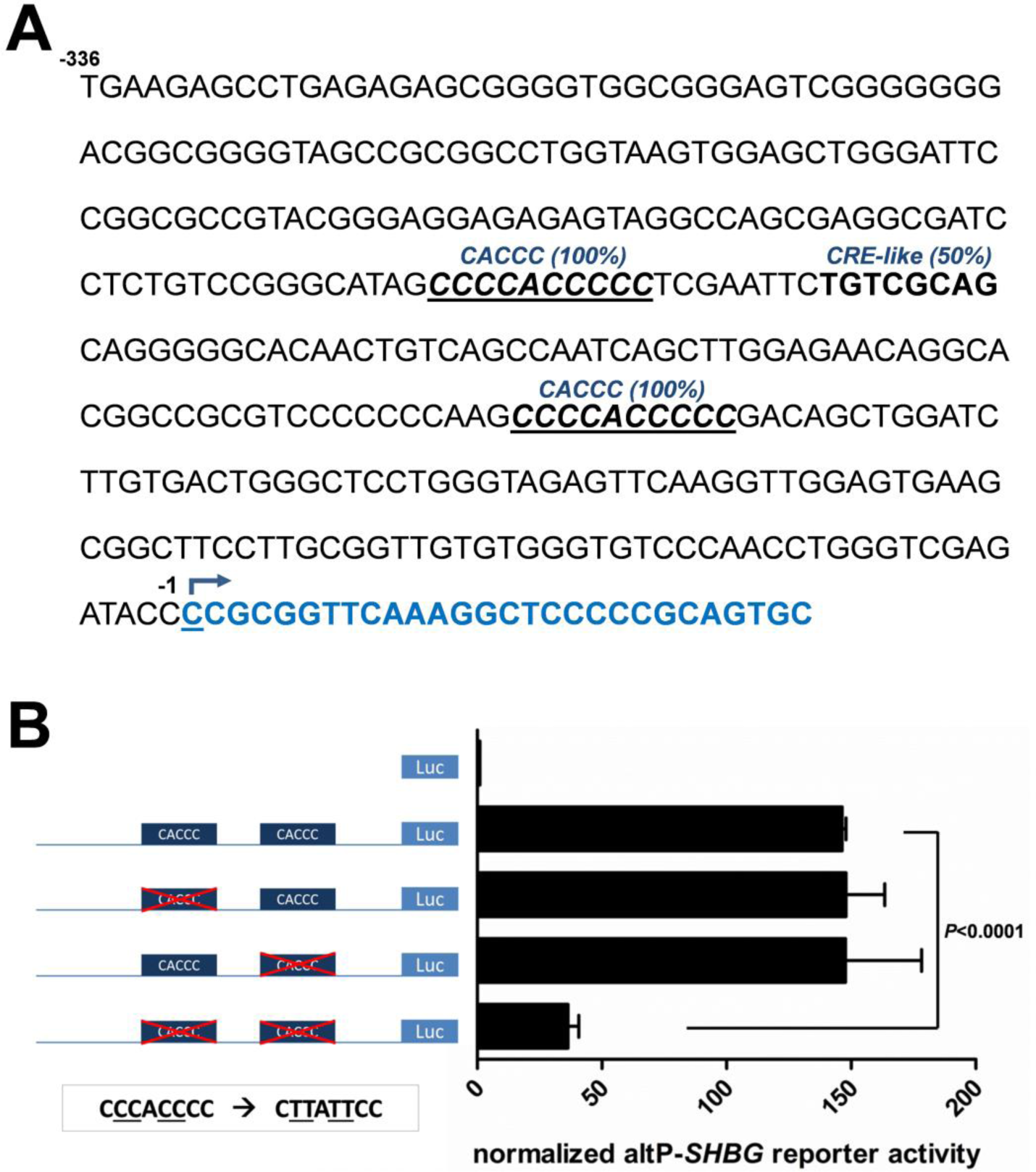
Dependence of altP-SHBG transcriptional activity on CACCC elements. A, Sequence analysis revealed two CACCC elements (underlined and bolded) and one partial CRE (bolded) within the minimal region of the alternative *SHBG* promoter. The transcriptional start site from this region (3) is also shown (underlined and arrow). Bases shown in blue are the first 28 nucleotides of exon 1A. B, The basal luciferase activities of altP-*SHBG* reporter constructs with single or double mutated CACCC elements were assessed in GC-2 cells. Relative luminescence counts are normalized to a promoterless pGL3 basic reporter vector that was assayed in parallel. Mutagenesis strategy is indicated. Data points are means ± SD of triplicate measurements. *P*-values and statistical significance were determined by one-way ANOVA.

Three mutant constructs were generated by site-directed mutagenesis using the wild-type construct as template. The distal, proximal or both of the CACCC elements were altered to be poor in GC content. These were transfected in parallel into GC-2 cells and their reporter activities were compared to the wild-type sequence. Figure 3B shows that while no changes in reporter activity are observed when only one CACCC element is mutated, disruption of both elements reduces this activity to ∼25% of wild-type (*P*<0.0001). I hypothesized that an endogenous factor within GC-2 cells must bind to either of these elements and drive optimal transactivation of the reporter gene.

### 2.2 Expression and localization of *Klf4* in GC-2 cells

The Krüppel-like family of transcription factors are also well known to interact directly with CACCC elements (18,19). Previous studies have demonstrated that *Klf4* is expressed during mouse spermatogenesis in round spermatids (20,21). This temporal pattern of expression bears striking similarity to when *SHBG* transcripts rise during the spermatogenic cycle and when SHBG is first observed in stage VII round spermatids within the 11K-SHBG transgenic mice (9,12). RT-PCR was performed on GC-2 cell cDNA using primers positioned at the 5’ and 3’ UTRs of the *Klf4* gene. Figure 4A shows detection of the full length 1613 bp *Klf4* open reading frame as previously described in the mouse testis (22), as well as several possible splice variants. Western blots performed on separate nuclear and cytoplasmic fractions from GC-2 cells demonstrated that KLF4 is confined within the nucleus (Figure 4B). Using a common forward primer positioned at exon 3 and unique reverse primers positioned at the four known polyadenylation signals (PAS) along the *Klf4* 3’UTR I identified usage of four PAS used by *Klf4* transcripts (Figure 4C).

**Figure 4.**
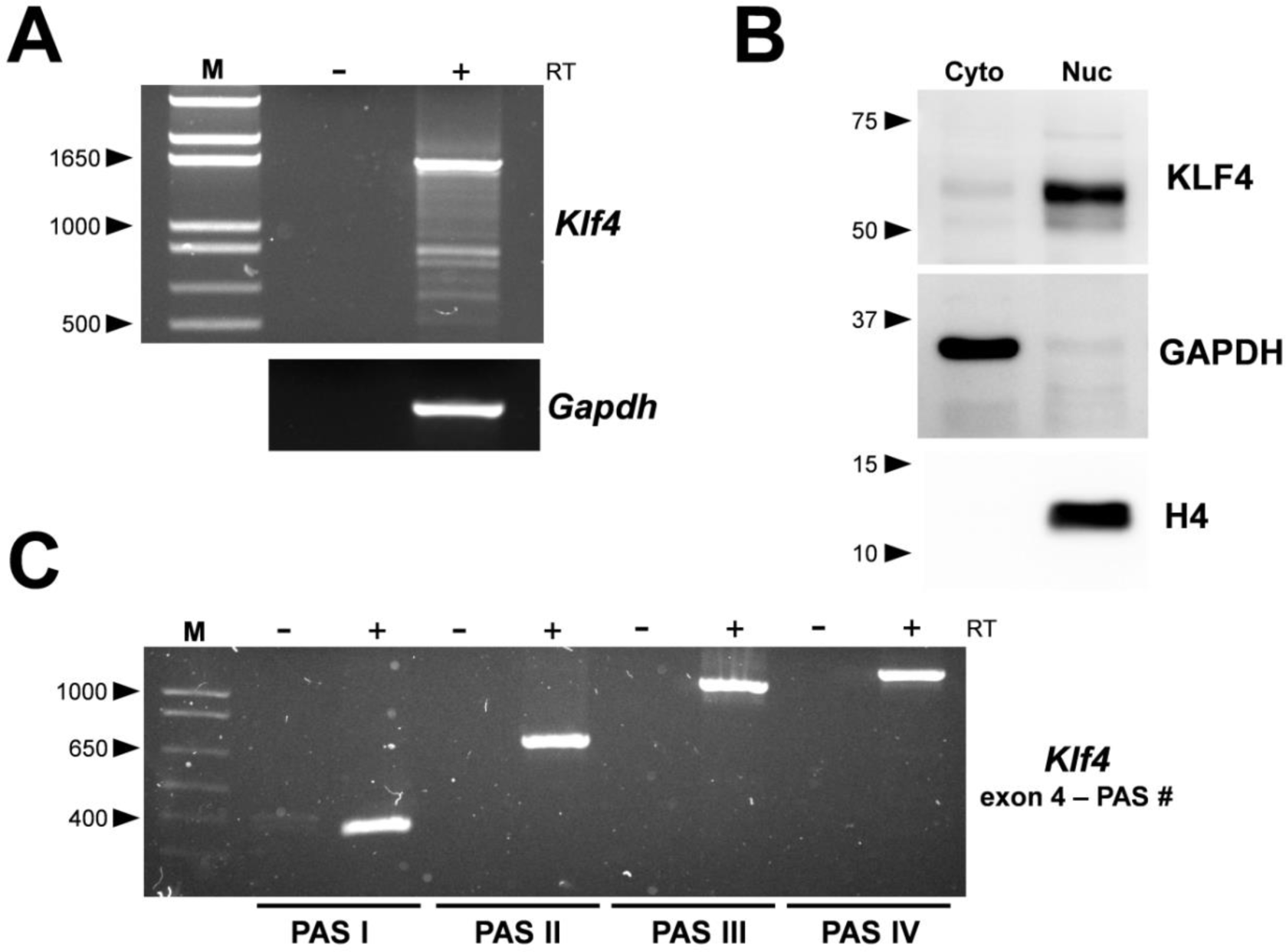
Expression and localization of *Klf4* in GC-2 cells. A, Full length mouse *Klf4* transcripts are detected in GC-2 cells by RT-PCR and electrophoresis in a 1% agarose gel. Molecular size in bp is indicated. Integrity of the cDNA in the sample is assessed by analysis for *Gapdh* transcripts. B, Western blot for KLF4 are performed on GC-2 cell cytoplasmic and nuclear fractions. Blots for GAPDH and histone H4 were performed on the same membrane to confirm the identity of the cytoplasmic and nuclear fractions, respectively. Molecular mass in kDa is indicated. C, Identification of four *Klf4* PAS utilized by GC-2 cells by RT-PCR and electrophoresis in a 1% agarose gel. Molecular size in bp is indicated.

### 2.3 High KLF4 levels are required for optimal altP-*SHBG* reporter activity in GC-2 cells

To test the impact of reduced KLF4 protein levels on altP-*SHBG* transcriptional activity, GC-2 cells were cotransfected with an siRNA targeting *Klf4* (si-*Klf4*) and the altP-*SHBG* reporter construct. Figure 5A shows that a decrease in KLF4 protein levels significantly reduces the activity of altP-*SHBG* compared to the control siRNA (*P*<0.001). Moreover, cotransfection of a *Klf4* overexpression construct with si-*Klf4* rescued the effects of knockdown compared to cotransfection with an empty vector (pcDNA) (Figure 5B).

**Figure 5.**
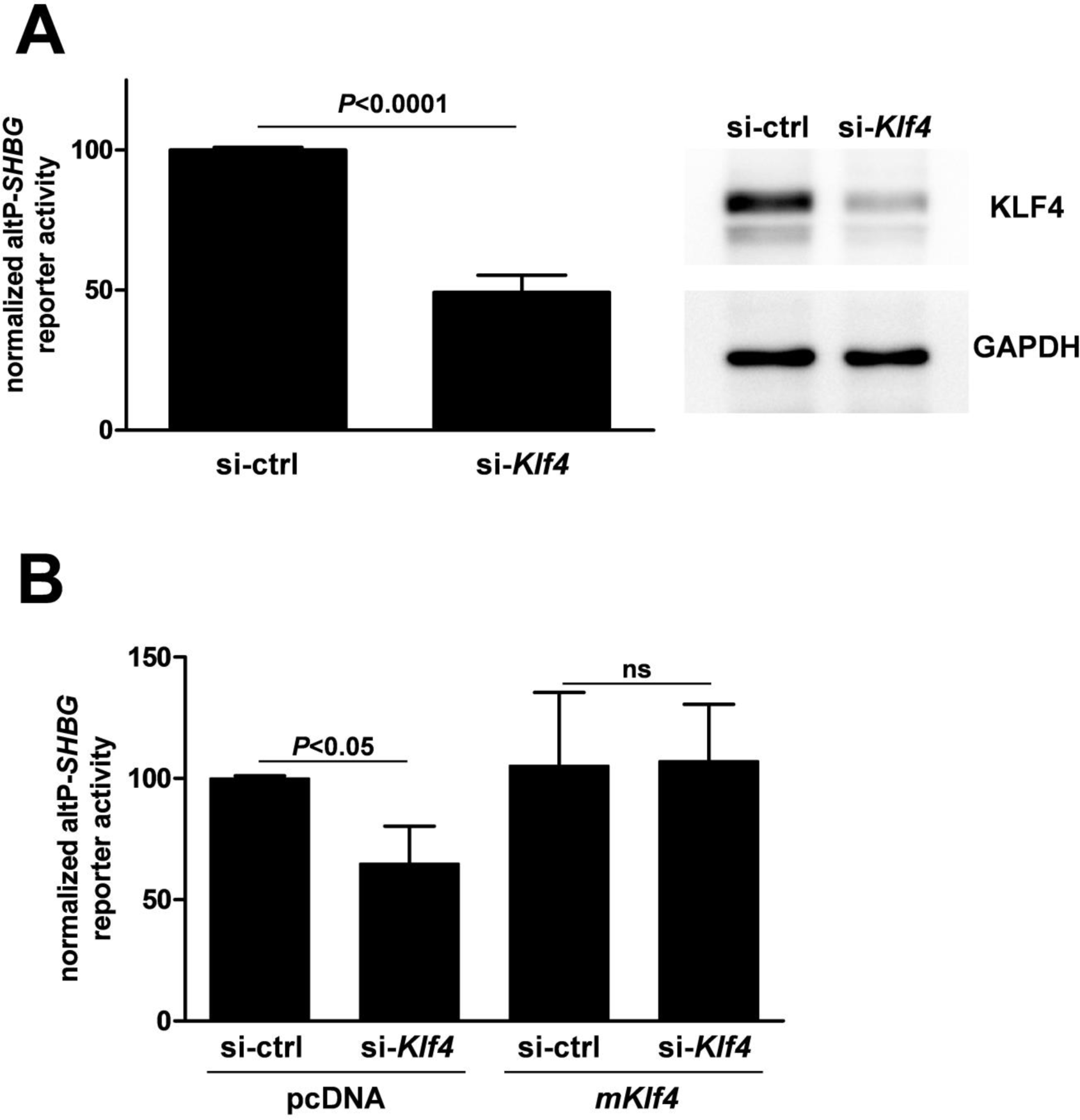
High KLF4 levels are required for optimal altP-SHBG reporter activity in GC-2 cells. A, The effect of reduced KLF4 levels on altP-*SHBG* reporter activity was assessed in GC-2 cells by cotransfection of the altP-*SHBG* reporter construct with either an siRNA targeting *Klf4* (si-*Klf4*) or non-targeting control (si-ctrl). Confirmation of knockdown was assessed by western blots for KLF4 on replicate GC-2 cell lysates from the same experiment. B, The effect of combined *Klf4* knockdown and overexpression on altP-*SHBG* reporter activity was assessed in GC-2 cells. Replicate knockdown experiments in A were cotransfected with either a *Klf4* overexpression construct or empty vector (pcDNA). For both A and B data points are means ± SD of triplicate measurements from three independent experiments. Data are normalized to the first mean in each histogram. *P*-values and statistical significance were determined by Student’s *t*-test. ns, not significant.

### 2.4 KLF4 localizes to the alternative *SHBG* promoter *in vivo*

To test the interaction of KLF4 with the alternative *SHBG* promoter *in vivo*, chromatin was isolated from the seminiferous epithelium of transgenic mice harboring the 11kb *SHBG* locus and subjected to probe sonication. Fourteen shear cycles yielded an optimal bolus of 200-300bp sized chromatin fragments (Figure 6A). This sample was used for chromatin immunoprecipitation (ChIP) using an antibody against KLF4 followed by PCR amplification of the alternative *SHBG* promoter (-250/-73) (Figure 6B). Chromatin integrity was assessed by ChIP with antibodies against RNAPII followed by PCR amplification of the *Gapdh* promoter.

**Figure 6.**
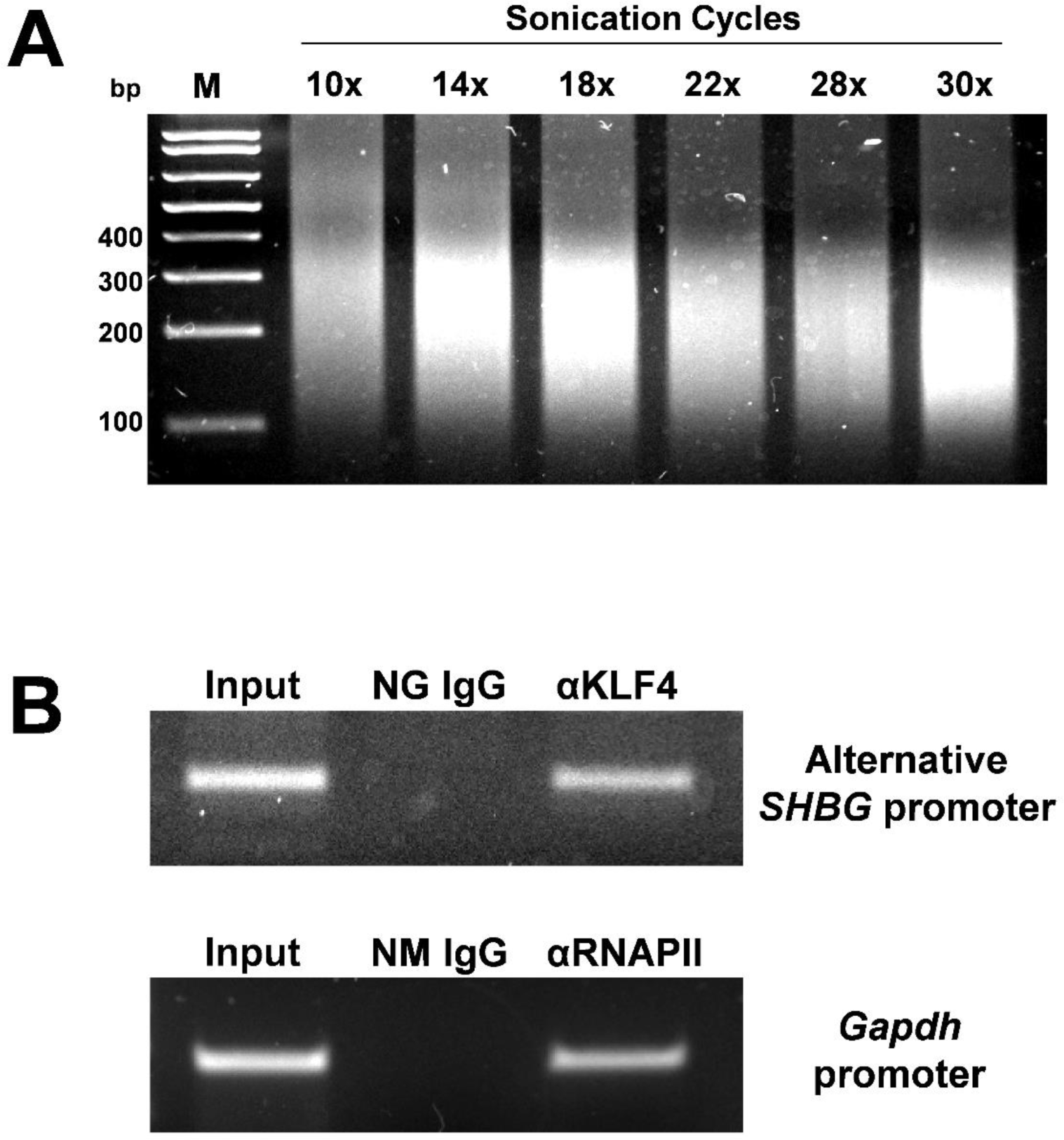
KLF4 localizes to the alternative *SHBG* promoter *in vivo*. A, Total chromatin from seminiferous tubules obtained from adult 11K-SHBG transgenic mice was fragmented by probe sonication. Sheared chromatin was reverse-crosslinked, purified and subjected to 2% agarose gel electrophoresis. B, ChIP with antibodies against KLF4 followed by PCR amplification of the alternative *SHBG* promoter (-250/-73). Chromatin integrity was assessed by IP with antibodies against RNAPII followed by PCR amplification of the *Gapdh* promoter. Input conditions were IPs performed with 1% of the chromatin reserved from IP with specific antibodies. PCR products were analyzed on a 1.5% agarose gel. Both gels were stained with SYBR. Molecular sizes in bp are indicated. M, molecular size marker.

## 3. Discussion

After meiosis, haploid testicular germ cells undergo a massive wave of transcription that results in the expression of numerous genes required for spermiogenesis (23,24). The rise in *SHBG* mRNA that occurs in stages VI-VII round spermatids (12) are likely part of this wave and their tightly regulated cyclical expression pattern suggests that mechanisms controlling their production are also spatiotemporally regulated. Round spermatids *in vivo* produce a specific environment of factors and mechanisms that permit transcription from exon 1A on the *SHBG* gene. GC-2 cells recapitulate this feature of round spermatids with high transcriptional activity from the alternative promoter while failing to produce any signal from the proximal *SHBG* promoter. Basal altP-*SHBG* reporter activity requires one intact CACCC element. Therefore the presence of two CACCC elements appears to be functionally redundant in this reporter model. KLF4 plays contrasting roles in different cellular contexts. It is required for the proper formation of the skin barrier as well as goblet cell differentiation in the digestive tract (25,26), yet is also one of the four Yamanaka factors that can induce pluripotency when introduced with Oct4/Pou5f1, Sox2 and c-Myc (27). In differentiating sperm, KLF4 is transiently but highly expressed in mid-to-late round spermatids in both the mouse and humans with transcript and protein levels undetectable by the time the cells are elongated (20,21,28). This transient expression pattern suggests that KLF4 plays a role in regulating post-meiotic gene expression.

Surprisingly, mice with *Klf4* ablated from their germ cells are completely fertile with no aberrations in spermatogenesis, nor differences in litter sizes sired from knockout males (21). Despite being dispensable for spermatogenesis, significant disturbances in testicular gene expression resulted in the mutant testes. By microarray analysis, the authors found 75 probe sets exhibiting downregulation and 90 probe sets exhibiting upregulation compared to wild-type, illustrating that KLF4 plays dual roles as both a transcriptional activator and repressor in the testis.

In GC-2 cells KLF4 localizes to the nucleus, consistent with its location *in vivo* in both mouse and human round spermatids (20,21,28). In the mouse testis, *Klf4* transcripts utilize four polyadenylation signals (PAS) (22). Sertoli cells utilize only the first PAS, while the latter three are considered markers for the presence of post-meiotic germ cells in the mouse testis. The observed pattern of *Klf4* expression and localization in GC-2 cells fully resembles its pattern in round spermatids *in vivo* and therefore supports the use of this cell line as a tool to model testicular germ cell expression of *Klf4*.

Transcriptional activity from the alternative *SHBG* promoter is significantly reduced when KLF4 levels are depleted by knockdown. While *Klf4* overexpression restored the effects of knockdown it did not consistently augment reporter activities above the control levels. This suggests that the reporter construct is fully saturated with KLF4. Nevertheless these findings support that KLF4 is a key transcriptional activator of *SHBG* in testicular germ cells. In differentiating stem cells KLF4 has been identified as a pioneer transcription factor that can access regions of the genome that are otherwise inaccessible due to ‘closed-state’ higher order chromatin structures (heterochromatin) (29,30). My ChIP analyses shows that KLF4 occupies the alternative *SHBG* promoter *in vivo*. Upon engaging exposed motifs on heterochromatin, pioneer factors are thought to further recruit chromatin modifiers, remodelers and other transcriptional coregulators to render these regions permissive for transcription (31,32). In support of this model, KLF4 has been shown to recruit the p300 coactivator to the promoters of KLF4-target genes, where it acetylates nearby histone tails to promote chromatin decondensation (33,34). The bridging function of KLF4 between promoter elements and transcriptional coregulators implies that it is part of a larger combination of factors that control *SHBG* expression. Nearly 50% of altP-*SHBG* reporter activity remains following *Klf4* knockdown and approximately 25% remains when both CACCC elements are disrupted suggesting that there may be other factors and elements at work.

Identification of a partial CRE element did raise my curiosity of whether germ cell *SHBG* is regulated by CREM, an essential transcription factor for mammalian germ cell development (35). My altP-*SHBG* reporter construct shows strong activation upon dibutyryl cyclic-AMP treatment and by overexpression of *CREM* isoforms however these effects are preserved upon mutation of the CRE element (Meyers & Hammond, unpublished), implying that these effects may be acting indirectly. The specificity protein (SP) family of transcription factors are wellknown to interact with and drive transcription from CACCC motifs (18,19). However, immunolocalization of SP1 and SP3 protein levels are only found in outer spermatogonial cells of the mouse testis (36) where *SHBG* is not expressed.

Transcriptional activation of *SHBG* by KLF4 is further supported by their shared expression patterns during the seminiferous cycle. Early characterization of 11K-SHBG transgenic mice revealed a stage-dependent expression pattern of *SHBG* transcripts during the seminiferous cycle (12). Intriguingly, peak levels of *Klf4* transcripts are also found in round spermatids at stage VII (20) which is also when *SHBG* transcripts reach their maximum. During post-natal development, *SHBG* transcripts in 11K-SHBG mouse testes begin to rise steadily after day 20 which is coincident with the first appearance of post-meiotic germ cells (37,38) and is also when *Klf4* mRNA dramatically increases (20).

## 4. Experimental Procedures

### 4.1 Animals

Mice expressing human *SHBG* transgenes (C57BL/6 X CBA background) (12) and their wild type littermates were maintained under standard conditions at the UBC Centre for Disease Modeling with food and water provided *ad libitum*. Tissues from adult male mice for chromatin immunoprecipitation were obtained under a protocol approved by the University of British Columbia Animal Care Committee.

### 4.2 Cell lines and transient transfections

Immortalized mouse spermatocyte cells, GC-2spc(ts) (catalogue number CRL-2196)(13) (GC-2) were obtained from the American Type Culture Collection (ATCC). Cells were maintained in Dulbecco’s Modified Eagle Medium, supplemented with 10% fetal bovine serum (FBS) and penicillin-streptomycin (100U/mL). Cells were grown in a standard laboratory cell culture incubator at 37°C in 100% humidity, 5% CO2 environment. Cells were routinely passaged 1:5 or 1:6 by detaching with trypsin-EDTA, twice weekly. To minimize the variable effects of cell line drift all experiments performed were with passage numbers below or equal to 15. Unless otherwise indicated, all cell culture reagents were purchased from Thermo Fisher Scientific.

### 4.3 Plasmids and reporter constructs

The alternative promoter fragment corresponding to -366/+28 nt with respect to the transcriptional start site at exon 1A was PCR amplified from a cosmid containing a fragment of the human chromosome 17 that was used previously for earlier sequencing studies of the *SHBG* gene (5). This fragment was cloned into the pGL3 luciferase reporter vector (Promega) by standard cloning methods. The primers used for sticky-end (XhoI/HindIII) cloning are listed in Table 1. The -266/+366 region of the proximal human *SHBG* promoter incorporated into pGL3 was constructed by Dr. Tsung-Sheng Wu as described (14). Within the promoter fragment, a stop codon was introduced immediately downstream of the translation initiation codon in exon 1. This was done to disrupt translation from the normal *SHBG* translation start site in exon 1 to ensure efficient translation initiation of the luciferase gene inserted within exon 2. The mouse *Klf4* construct (within pcDNA 3.1) was a generous gift from the laboratory of Dr. J.P. Katz via Dr. Marie-Pier Tetreault at the University of Pennsylvania, Philadelphia, USA.

**Table 1.**
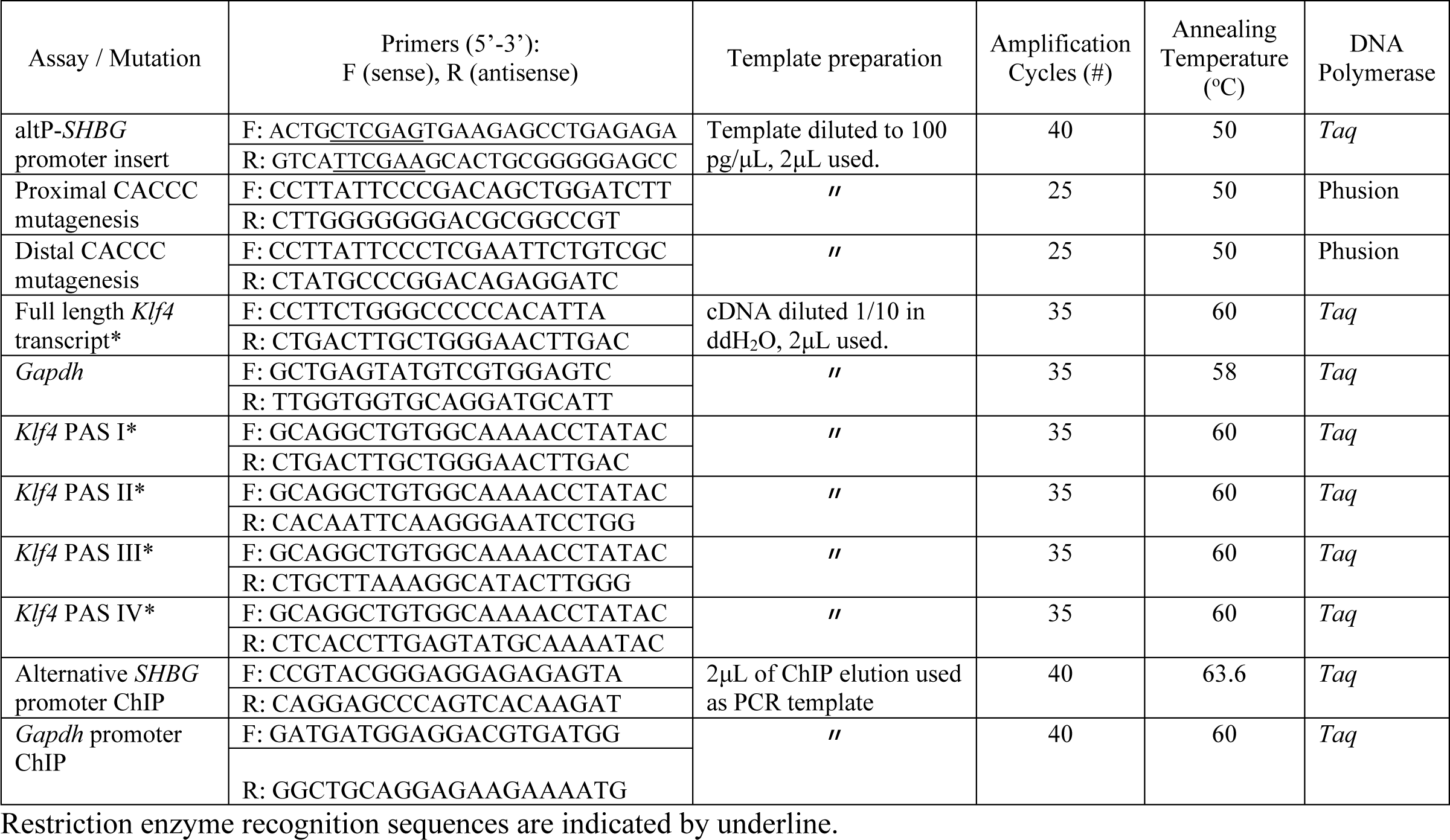
Oligonucleotide sequences and PCR parameters used in this study.

### 4.4 Transient transfection and luciferase reporter assay

Approximately 16-18 hours before transfection, GC-2 cells were trypsinized from maintenance culture into a single cell suspension, counted and 1.4x10^5^ cells/well were seeded into 24 well plates in 0.5mL of culture media. The next morning each well was changed and transfections were performed as follows. Transient cotransfection of luciferase and β-galactosidase reporter constructs were performed using Lipofectamine 2000 transfection reagent according to the manufacturer’s protocol. In all experiments a ratio of 2µL of transfection reagent to 1µg of nucleic acid was used. After separately diluting the transfection reagent and pool of nucleic acids in Opti-MEM, the two dilutions were combined and incubated for 30min before pipetting onto cell media. Each well of cells was transfected with 500ng of luciferase reporter construct plus 45ng of pRC-CMV-*LacZ* (β-galactosidase) unless otherwise indicated in figure captions. For experiments involving cotransfection with additional plasmids or siRNA, the transfection complex incubation time was increased to 45-60min before pipetting them onto cell media.

Media was refreshed 6-7hrs after transfection.

Forty-eight hours after transfection the cells were washed once with 0.5mL sterile PBS and 75uL of 1x Reporter Lysis Buffer (Promega) supplemented with protease inhibitor cocktail lacking EDTA (Thermo Fisher Scientific) was pipetted onto each well. Cells were allowed to swell for 5min at room temperature before subjecting them to a single freeze/thaw cycle at -80°C. Crude lysates were harvested from their wells by pipetting and transferred to separate microcentrifuge tubes. These were centrifuged at 18,000g for 10min at 4°C before transferring supernatants to fresh tubes. Separate 25µL aliquots of each lysate were plated onto white opaque and clear bottom 96 well plates for measurement of luciferase and β-galactosidase activities, respectively.

Using a multichannel pipet, 65µL of Luciferase Assay Reagent (Promega) was added in unison to the lysates and luminescence activity in counts per second were immediately recorded on a Victor X4 plate reader (Perkin Elmer). Using a multichannel pipet, 35µL of 1x β-galactosidase assay buffer (Promega) was added in unison to plated lysates and mixtures were incubated at 37°C for 40 min. Reactions were stopped by adding 75uL of 1 M sodium carbonate and absorbance at 405nm (A405) was immediately recorded on a Victor X4 plate reader. The luminescence activity value of each well was divided by its respective A405 reading to correct for transfection efficiency to obtain a relative luminescence value.

### 4.5 Transcription factor binding site and CpG island analysis

Two online tools were used to screen for putative transcription factor binding sites: TRANSFAC (http://www.gene-regulation.com/pub/databases.html) and ORKAtk (http://cisreg.cmmt.ubc.ca/cgi-bin/ORCAtk/orca). Searches using both programs were performed with a cutoff threshold of 80% match to consensus binding sites. Only hits obtained from both analysis tools were considered as putative binding sites. The CpG Islands locator tool (http://www.bioinformatics.org/sms2/cpg_islands.html) was used for CpG island detection.

### 4.6 Small interfering RNA (siRNA)

Knockdown of mouse *Klf4* transcripts was performed with an ON-TARGETplus SMARTpool siRNA against mouse *Klf4* (L-040001-01-005, 16600) (Dharmacon, GE Healthcare). This pool of four targeting RNAs targets the following sequences: AGAUUAAGCAAGAGGCGGU, CCAUUAUUGUGUCGGAGGA, CCGAGGAGUUCAACGACCU,CGACUAACCGUUGGCCGUGA. Negative control conditions were performed using an ON-TARGETplus Non-targeting Control siRNA from the same manufacturer.

### 4.7 RNA extraction and cDNA synthesis

Total RNA was extracted from cells or tissues using the RNeasy Mini Kit (Qiagen, Toronto, Ontario) with a mid-protocol DNase I treatment. Concentrations of eluted RNA were determined using a NanoDrop 2000 spectrophotometer (Thermo Fisher Scientific). Unless used immediately, RNA samples were stored at -80°C. Total RNA (2500ng) was reverse transcribed using SuperScript® II Reverse Transcriptase with oligo(dT) primers (Thermo Fisher Scientific) in a 20uL volume according to the manufacturer’s instructions with the exception that the synthesis incubation period at 42°C was increased from 50 to 60min. All cDNA samples were stored at - 20°C.

### 4.8 Primers and polymerase chain reaction (PCR)

Oligonucleotide/primer sequences for all PCR reactions described in this thesis are listed in Table 2.1. Amplification reactions were performed with either PCR Supermix (Thermo Fisher Scientific, catalogue no. 10572014) containing *Taq* polymerase, Phusion^TM^ High-Fidelity DNA polymerase for mutagenesis of C-G rich templates (Thermo Fisher Scientific, catalogue no. F541) or AccuPrime^TM^ *Pfx* DNA polymerase (Thermo Fisher Scientific, catalogue no. 12344024), each with their included reaction buffers. All PCR reactions were performed in 25μL reaction volumes with primers diluted according to the instructions included with each polymerase. All reactions were carried out in a iCycler thermocycler (Bio-Rad) with the lid held at 100°C. Whole reaction volumes were analyzed by agarose gel electrophoresis stained with SYBR® (catalogue no. S33102) diluted 1:10 000. Gel images were captured on a ImageQuant LAS 4000 system (GE Healthcare Life Sciences)

### 4.9 Whole cell extracts and subcellular fractionation

GC-2 cells were lysed by re-suspension in 1x Cell Lysis Buffer (Cell Signalling Technology, Danvers, MA) supplemented with protease inhibitor cocktail (Thermo Fisher Scientific, Waltham, MA) and 1mM phenylmethane sulphonyl fluoride (PMSF) followed by a single freeze-thaw cycle according the manufacturer’s protocol. Preparation of nuclear and cytoplasmic fractions was performed using the NE-PER Nuclear Protein Extraction Kit (Thermo Fisher Scientific, Waltham, MA) according to the manufacturer’s included protocol. The total protein content of all lysates was quantified in a DC (detergent compatible) Protein Assay (Bio-Rad) against a standard curve of BSA diluted in lysis buffer according to the manufacturer’s instructions.

### 4.10 Seminiferous tubule isolation from mouse testes

Following isoflurane overdose, male mice were perfused with room temperature PBS for 10- 15min until the perfusate from the animal was clear. The testes were quickly dissected out and placed in ice-cold DMEM for 5-10min. Each testis was transferred to a dish containing ice cold PBS, decapsulated and tubules were placed into a fresh sterile tube containing 7.5mL of DMEM with 0.8mg/mL collagenase type 1A, 0.225mg/mL hyaluronidase, 0.3mg/mL DNase I (digest media). Tubes were incubated at 37°C shaking gently and were gently flicked every 5min to separate seminiferous tubules from the interstitial cells and material. After 20min, tubes were removed from the incubator and tubules were allowed to sediment by gravity. Supernatants were removed and tubules were resuspended in another 7.5mL of fresh digest media before returning to 37°C for an additional 10-15min. Tubules were allowed to sediment again and supernatants were removed. Tubules were then washed four times by re-suspending in 9-10mL PBS and centrifuging at 180g for 5min.

### 4.11 Seminiferous tubule isolation for and chromatin immunoprecipitation

Seminiferous tubules from three 11K-SHBG transgenic mouse testes were isolated as described above and pooled together. Tubules were fixed in freshly prepared 1% paraformaldehyde dissolved in 1xPBS, pH 7.4 for 10min at room temperature. Tissue lysis and chromatin isolation was performed with a Magna ChIP^TM^ A/G kit (Millipore) according to the manufacturer’s protocol with the exception that following lysis, crude lysates were passed 14 times through a 22 gauge needle fitted to a sterile 5mL syringe. Chromatin-containing supernatant (600μL) was divided into three 200μL aliquots and stored at -80°C.

All of the following steps for chromatin shearing were performed on ice using ice-cold buffers and tubes. One testis’ worth of chromatin (200μL) was resuspended in 1500μL of ice cold Shearing Buffer (Millipore) supplemented with EDTA-free protease inhibitor cocktail (Thermo Fisher Scientific) and 1mM PMSF and aliquoted into six 255uL in separate 1.5mL tubes.

Chromatin was sheared using a Microson Ultrasonic Cell Disrupter XL probe sonicator while on ice. The optimal condition to shear all chromatin into 200-300 bp fragments was 14 cycles of: 5s sonication pulse followed by 25s rest to prevent overheating. Sample tubes were immersed in an ice bath the entire time. Sheared samples were centrifuged at 18,000g for 10min at 4°C and supernatants were aliquoted into four 50μL chilled microcentrifuge tubes and stored at -80°C for subsequent ChIP assays. Remaining volumes (∼55μL) were reverse cross-linked as described in the provided kit protocol (Millipore) by incubating at 62°C overnight in a kit-supplied ChIP Elution buffer supplemented with 0.33 µg/µL proteinase K (Thermo Fisher Scientific). The next morning DNA fragments were cleaned up using a PCR Purification Kit (Qiagen) and eluted in 30uL. This volume was subjected to 2% agarose gel electrophoresis and visualized as described above to confirm shearing efficiency. ChIP analysis was performed using the Magna ChIP^TM^ A/G kit (Millipore) following the included protocol exactly as described with the KLF4 antibody below. Oligonucleotide primers for PCR analysis are listed in Table 1.

### 4.12 Polyacrylamide gel electrophoresis (PAGE) and western blotting

Whole cell lysates or nuclear and cytoplasmic fractions (22μg) were resolved by 10% or 12% SDS-PAGE and electro-transferred to 0.45μm PVDF mini membranes using a Trans-Blot Turbo Transfer system (Bio-Rad). Gel to membrane transfers were performed at 25V, 1.3A for 10min. The membranes were blocked in 1x PBS, 0.1% tween 20 (PBS-T) containing 5% skim milk (blocking buffer) for 1hr at room temperature, followed by overnight incubation at 4°C with primary antibodies diluted in blocking buffer. The next morning membranes were washed once with blocking buffer for 5-10min and incubated with horseradish peroxidase-conjugated secondary antibodies (Sigma-Aldrich) diluted 1:10 000 in blocking buffer. Immunoreactive proteins were detected in a ImageQuant LAS 4000 system with Amersham ECL Prime chemiluminescent reagent according to the manufacturer’s protocol (GE Healthcare Life Sciences). Primary antibodies used in this study were KLF4 (AF3158) from R&D Systems, GAPDH (V-18, sc-20357) and histone H4 (Lys 8-R, sc-8660-R) from Santa Cruz Biotechnology.

